# Degradation of rutin and genistein and the effect on human bacterial fecal populations of obese and non-obese

**DOI:** 10.1101/2025.03.16.643526

**Authors:** Julia Jensen-Kroll, Tobias Demetrowitsch, Sabrina Sprotte, Fynn Brix, Alexia Beckmann, Kristina Schlicht, Erik Brinks, Matthias Laudes, Silvio Waschina, Charles M.A.P. Franz, Karin Schwarz

## Abstract

This study re-evaluates the microbial degradation of rutin and genistein, emphasizing health-related metabotypes and novel degradation and transformation products of these compounds. *In silico* predictions and calculations were made to determine the molecular formulae of metabolites derived from precursor molecules by methylation, sulfation, and dehydroxylation. To validate these, anaerobic, *ex vivo* experiments were conducted with pooled fecal samples of obese and non-obese test persons (n=7 per group) over 48 hours, followed by 16S rRNA gene sequencing and semi-targeted high-resolution mass spectrometry analysis. We identified 46 rutin and 23 genistein metabolites, including novel methylated and sulfated derivatives. Microbial analysis revealed that the plant compounds influenced 34 bacterial families and 83 genera. Rutin inhibited obesity-associated genera while promoting butyrate-producing bacteria in BMI >40 samples, but reduced health-associated bifidobacteria and increased enterobacteria. Both compounds stimulated *Eggerthella.* These findings highlight significant microbial and metabolic shifts induced by plant-derived compounds, suggesting potential health implications.

## 1 INTRODUCTION

Polyphenols are the largest and most diverse group of plant compounds. With over 8000 compounds, they are of great importance in human nutrition (Ross & Kasum, 2002). They are naturally present in fruits, vegetables, and grains, as well as in beverages such as tea, coffee, juices, and wine, and exhibit numerous beneficial properties, including antioxidant, anti-inflammatory, anticancer, immunomodulatory, and gut microbiome-modulating effects. The latter effect is partly due to the fact that polyphenols have low bioavailability and largely reach the colon undigested, where they can interact with the associated microbiota (Cardona et al., 2013). On the other hand, it is often not the polyphenolic precursor itself, but rather the microbial degradation product that is associated with the health-promoting effect or bioactivity. For example, Najmanová et al. (2016) demonstrated in a rat model that 3-(3-hydroxyphenyl) propionic acid, a microbial degradation product of flavonoids, can lower arterial blood pressure. Moreover, Lin et al. (2007) showed in an *in vitro* cell model that 3,4-dihydroxybenzoic acid, a degradation product of various polyphenols (Mosele et al., 2015), induces apoptosis in gastrointestinal cancer cells through the MAPK (mitogen-activated protein kinase) cascade.

In addition to the breakdown of dietary components, the healthy human microbiome has numerous other functions. These include the synthesis of vitamins and essential amino acids, the production of short-chain fatty acids (SCFAs), the metabolism of drugs and xenobiotics, the maintenance of the intestinal barrier, and the protection against pathogens (Jandhyala et al., 2015).

The microbial community is influenced by several factors, including diet, lifestyle, age, genetics, and diseases (Gomaa, 2020). When the balance of the microbial community is disturbed, it is referred to as dysbiosis or a dysbiotic microbiome. Several diseases have been associated with dysbiosis, i.e., inflammatory diseases such as inflammatory bowel disease and ulcerative colitis, neurodegenerative diseases such as Parkinson’s disease, and metabolic disorders like coronary heart diseases, obesity, or adiposity (Gebrayel et al., 2022).

In a systematic review of various studies, Pinart et al. (2021) identified 15 bacterial genera that appear to be dysregulated in the context of obesity when compared to a healthy microbial community. This provided additional markers besides a generally less diverse microbiota and the controversially discussed altered Firmicutes to Bacteroidetes (F:B) ratio. Obesity is also characterized by an excessive accumulation of body fat, often resulting from a combination of genetic, environmental, and lifestyle factors (Romieu et al., 2017). It has received considerable attention in recent years, as it can lead to a wide range of health complications, including cardiovascular diseases (CVD), type 2 diabetes (T2D), non-alcoholic fatty liver disease (NAFLD), and respiratory problems (Geng et al., 2022). The debate as to whether dysbiosis is the cause, or a consequence of the disease, is still ongoing. The first associations between the gut microbiota and obesity were described by Turnbaugh et al. (2006). They reported that some microorganisms can extract more energy from food and make it available to the host. In addition, the study by Tims et al. (2013) showed that obesity is associated with an increase in carbohydrate-fermenting bacteria and SCFA producers. As reviewed by Sanmiguel et al. (2015), saccharolytic bacteria play a dual role in human health. They are essential for digestion, by fermenting carbohydrates to produce short-chain fatty acids (SCFAs) such as acetate, propionate, and butyrate, which can serve as energy sources for the host. However, excessive production of these metabolites can increase energy expenditure, particularly in overweight individuals, potentially increasing the risk of obesity. A critical factor influencing this dynamic is the F:B ratio. This is because Firmicutes are particularly efficient at extracting energy from food, leading to higher caloric intake. In addition, saccharolytic bacteria influence metabolism by regulating glucose homeostasis, modulating insulin sensitivity, and stimulating satiety hormones such as glucagon-like peptide-1 (GLP-1) and peptide YY (PYY). Microbial imbalances can disrupt these pathways, making appetite regulation and weight control more difficult. In essence, while saccharolytic bacteria are indispensable for healthy digestion, their dysregulation or interplay with a high-caloric diet can significantly contribute to obesity as reviewed (Sanmiguel et al., 2015). Polyphenols have been shown to have a beneficial effect on obesity in numerous studies, as recently reviewed by Aloo et al. (2023). The main mechanisms reported are that polyphenols inhibit obesity through several processes, including inhibition of digestive enzymes (particularly alpha-glucosidase, pancreatic lipase, fatty acid synthase, and alpha-amylase), stimulation of energy expenditure, suppression of appetite, regulation of lipid synthesis, and modulation of the gut microbiota. Chen et al. (2024) showed in a mouse model that these effects may be mediated not only by the polyphenolic precursor molecules but also from the microbial degradation products. They showed that 3- (3’,4’-dihydroxyphenyl) propanoic acid and 3’,4’-dihydroxyphenylacetic acid successfully prevented obesity in mice on a high-fat diet by regulating key metabolites within lipid, amino acid, and carbohydrate pathways. However, many mechanisms and the response of the microbial community are not yet fully understood.

A significant need for research is also in the area of metabotypes, as highlighted in the review by Hu et al. (2024). In particular, there are significant gaps in the area of disease-related metabotypes. Metabotypes could be seen as markers of microbial activity and could be used to assess how healthy a microbial composition is on a scale from healthy to dysbiotic. Hu et al. (2024) specifically mentioned that especially the combination of metagenomics and metabolomics can significantly improve the understanding of (poly)phenolic metabolism and the role of the gut microbiota. In particular, metabolomics allows the assessment of functional aspects that are often overlooked when analyzing the composition of the gut microbiome (Hu et al., 2024).

Although there are numerous reports on rutin and genistein, in our opinion the topic of disease-related metabotypes is lacking. Quercetin including its glycosides and genistein play a central role in today’s diet. Quercetin, is the most widespread polyphenol in the plant kingdom, with approximately 128 known glycosides. Rutin, a glycosylated derivative of quercetin, enters the colon largely undigested. Genistein, an isoflavonoid, is commonly found in soy and soy-based products along with daidzein and glycitein. Soy products are often associated with healthy diets, such as vegetarian diets, which are gaining popularity among consumers, especially in the context of meat substitutes. However, in contrast to daidzein, there is limited knowledge about metabotypes for genistein (Hu et al., 2024; Soukup et al., 2023).

The present study, reconsidered rutin and genistein with a deeper focus on the health-related metabotypes and an entirely fresh look at the degradation and transformation products. In particular, the study of transformation products is, to our knowledge, underrepresented. Based on the assumption that bacteria are also capable of transformation processes such as methylations and sulfation, and considering that methylation, sulfation, and, for example, dehydroxylation can occur multiple times, molecular formulae for metabolites derived from precursor molecules were predicted and calculated *in silico*. To validate these predictions and to investigate a possible disease-related metabotype, *ex vivo* experiments were performed with pooled fecal samples from healthy subjects (BMI <25) and obese subjects (BMI >40). The work is based on the experience gained from the use of fecal sample in *in vitro* systems in general, as well as the pooling of fecal samples (Aguirre et al., 2014^a^; Aguirre et al., 2014^b^ Aguirre et al., 2015; Aguirre, et al., 2016; Aguirre et al., 2017; Pérez-Burillo et al., 2021). In addition, to analyze the presence of metabotypes and novel, previously undescribed degradation and transformation products, the study also examined how the microbial community changes in response to plant compounds and whether there are differences between the two BMI pools in this regard.

## 2 MATERIAL AND METHODS

### 2.1 Sample material

The human fecal samples originated from 2 different groups of volunteer test persons. One group of individuals (n=7) with a BMI >40 kg/m^2^ and one (n=7) with a BMI of 18.5-25 kg/m^2^. For both groups, an exclusion criterion was the intake of antibiotics within the last 3 months.

### 2.2 Ethical aspects

The volunteers were informed about the use of the sample material in advance of the study by the supervising clinician; no personal data were collected. The only data recorded concerned data on the BMI status. The basis for the collection and use of the sample material was stipulated by the ethics application with reference number AZ D 420/20 (Ethics Committee, Medical Faculty, Kiel University).

### 2.3 Sample collection

The fecal samples were transferred to a sample vessel immediately after donation and stored cooled and under anaerobic conditions in an anaerobic jar (AG0025A Oxoid, Thermo Fisher Scientific, Dreieich, Germany). An AnaeroGen^TM^ pack (Oxoid, Thermo Fischer Scientific, Dreieich, Germany) was added to the jar to generate an anaerobic atmosphere. The samples were transferred into an anaerobic workstation (Whitley A45, Meintrup DWS Laborgeräte, Herzlake, Germany) and aliquoted under anaerobic conditions within 24 h. Fecal samples were then stored at -80°C until the preparation of fecal cultures.

### 2.4 Anaerobic incubation

Rutin and genistein were purchased from Cayman Chemical (Tallinn, Estonia) and had a purity of ≥98%. These phytochemicals were dissolved in DMSO (SERVA, Heidelberg, Germany), filtered using a 0.2 µm disposable membrane filter (Filtropur S, Sarstedt, Nümbrecht, Germany), and kept at 4°C until use. The fecal samples of all subjects were transferred to the anaerobic workstation. The anaerobic gas mixture in the workstation consisted of 10% H2, 10% CO2, and 80% N2. First, the seven fecal samples from individuals of each of the two groups were pooled following the protocol of Pérez-Burillo et al. (2021) and the experiments of Aguirre et al. (2014^a^, 2014^b^, 2015, 2016, 2017). As it is well known, that the human gut microbiota can differ substantially between individuals, we opted to pool fecal samples from individuals in this study in order to maximize the diversity of microorganism associated with the characteristic microbiota of the BMI groups. Thus, an effect on any of the complete range of microbiota possibly present and associated with the specific BMI group, would be detected. The pooled fecal sample consisting of aliquots of 2.4 g of each of the seven samples was incubated in 315 ml (5.3% w/v) of modified BHI broth [37 g BHI dissolved in 1 l ddH2O, boiled, cooled with stirring at RT while flushing with 0.5 bar N2 for 30 min, followed by autoclaving]. 25 µg/ml L-cysteine monohydrochloride hydrate (Sigma-Aldrich, Taufkirchen, Germany), 5 µg/ml hemin (AppiChem, Darmstadt, Germany), and 0.1 µg/ml vitamin K1 (Carl Roth, Karlsruhe, Germany) were added, and the solution was transferred to a tinted pressure plus flask with a bromobutyryl septum (Duran®, Schott, Mainz, Germany)] for 1 h at 37°C with stirring (150 rpm; Mix 8 XL, 2mag AG, München, Germany) for dissolving and anaerobic adaption. For the exact composition of the BHI medium, **see supplement**. Afterward, six replicate samples with a ca. 1% fecal concentration (i.e., 10 ml mixed with 45 ml BHI) in modified BHI broth were prepared from each starter pool. The plant compounds were added to three of the replicate samples per group with a concentration of 0.5 mM. The other three replicate samples were used as a control. Two further controls, the stock solutions of the plant compounds in DMSO and the plant compounds in BHI broth, were subjected to the same cultivation conditions. The batch incubation process was performed in tinted flasks with bromobutyryl septum over 48 h. At 0, 6, 8, 24, and 48 h, a 2 ml sample was removed from each batch for metabolomic analysis. Samples were subsequently centrifuged (0.5 min, 12,700 *xg*), and the supernatant was filtered (0.2 µm, Filtropur S, Sarstedt, Nümbrecht, Germany) under a clean bench and stored at -80°C until extraction.

### 2.5 DNA isolation from feces and 16S rRNA gene metagenomic analysis

In parallel with the metabolomic analysis, a 16S rRNA gene metagenomic analysis was performed. For this, 1 ml was taken from each of the above samples at 0, 24, and 48 h and centrifuged for 2 min at 12,700 *xg*. The supernatants were discarded, and the pellets were frozen at -80°C until DNA purification. DNA was purified using the NucleoSpin DNA Stool Mini Kit (Macherey-Nagel, Düren, Germany) according to the manufacturer’s instructions. DNA concentration and purity were determined using a Qubit 3 fluorometer and a Nanodrop 2000c (Thermo Fisher Scientific, Dreieich, Germany), respectively.

For metagenomic analysis, the V3/V4 region of the 16S rRNA gene was amplified according to the 16S Metagenomic Sequencing Library Preparation Guide provided by Illumina (Illumina, Berlin, Germany), except for that Q5 polymerase was used (New England Biolabs, Frankfurt am Main, Germany). In brief, the library preparation consisted of a two-step PCR. In the first step, the target was amplified using the 2x Q5 Hot Start Polymerase Mix, while in the second PCR, the index barcodes were added to the amplicon using the 2x Q5 Ultra II MasterMix (New England Biolabs).

The concentration of the purified amplicons was measured using a Qubit 3.0 fluorometer (Invitrogen, Thermo Fisher Scientific, Schwerte, Germany). An average DNA fragment size of ∼ 580 bp was determined from 11 randomly selected samples using the Experion electrophoresis station with the DNA 1K Assay (Bio-Rad, Feldkirchen, Germany). Based on the data, the molarity of each sample was determined using the following formula: DNA concentration (ng/µL)/ [(660 g/mol x average DNA fragment size (bp)]x 10^6^.

The samples were diluted to a molarity of 4 nM using 10 mM Tris/HCl, pH 8.0, and pooled in an equimolar manner. The resulting pool was further prepared for loading on the MiSeq following the denaturation and dilution protocol from Illumina. The resulting library was diluted to a loading concentration of 10 pM, and a PhiX control was added to a final proportion of 5%. The final library pool, including the PhiX control, was sequenced using a MiSeq Reagent Kit v3 (600 cycles) and sequenced with 2x 301 cycles on a MiSeq high-throughput sequencer (Illumina, München, Germany).

### 2.6 Data handling and evaluation 16S rRNA gene metagenomics data

Primer sequences were removed from the reads using the software cutadapt version 3.7 (Martin, 2011). Sequencing reads in FASTQ format were analyzed using the R-package DADA2 version 1.20.0 (Callahan et al., 2016). Reads were filtered and trimmed using the function ‘filterAndTrim’ in its default parametrization except for the options ‘truncLen’ and ‘maxEE’. ‘truncLen’ was set to ‘c(255,235)’ to truncate forward reads to 255 bases and reverse reads to 235 bases. ‘maxEE’ was set to ‘c(2,3)’ to discard all sequences with an expected error number of >2 bases in forward reads and >3 bases in reverse reads after sequence trimming. Amplicon sequence variants (ASVs) were inferred using DADA2’s core function ‘dada’. Paired-end reads were merged using the function ‘mergePairs’ with a minimum sequence overlap of 14 bases, not allowing any mismatches. Unmerged reads were discarded from further analysis. Bimera ASV sequences were detected and removed using the function ‘removeBimeraDenovo’. The taxonomic classification of ASVs was predicted using the function ‘assignTaxonomy’ using the reference training data set from SILVA NR99 version 138.1 (Quast et al., 2013).

Alpha-diversity metrics were calculated using the R-package vegan version 2.6.4 (Oksanen et al., 2017) from the ASV count table. To estimate the pairwise beta-diversity of microbiota profiles, the ASV count table was first normalized by dividing counts by the per-sample Geometric Mean of Pairwise Ratios (GMPR, (Chen et al., 2018)). Student’s t-tests were used to assess the statistical significance of alpha-diversity between genistein/rutin-treated *ex vivo* cultures versus control cultures (no treatment) at the same times of sampling.

Based on the normalized ASV count table, pairwise Bray-Curtis dissimilarity scores were calculated as beta-diversity estimates. Classical multidimensional scaling (MDS) was applied for the results presented in **Fig. 2** using the R-function ‘cmdscale’ with k=2 to represent the beta-diversity data on two dimensions.

Differential abundance analysis was performed on family and genus taxonomic levels. To this end, ASV counts with identical family or genus taxonomic classification were aggregated to obtain family and genus count tables, respectively. Differential abundance of taxa between genistein/rutin-treated *ex vivo* cultures versus control cultures (no treatment) was tested at each time point (0 h, 24 h, 48 h) independently using the R-package DESeq2 version 1.40.1 (Love et al., 2014). Statistically significant differences in taxa relative abundance were assessed using Wald tests as implemented in DESeq2’s function ‘nbinomWaldTest’. As size factors, the Geometric Mean of Pairwise Ratios (GMPR) was calculated for each sample (Chen et al., 2018). P-values were adjusted for multiple testing using the method of Benjamini and Hochberg (1995).

### 2.7 Sample preparation for metabolomics analyses

For the extraction of hydrophilic and lipophilic compounds, 300 µl of each filtered supernatant was used. Samples were extracted using a modified SIMPLEX protocol according to Jensen-Kroll et al. (2022) (Jensen-Kroll et al., 2022). The samples were separated into proteinogenic, lipophilic, and hydrophilic fractions. The proteinogenic fractions were not further examined. The hydrophilic and lipophilic fractions were completely dried by centrifugal evaporation (SpeedVac concentrator (Savant SC250EXP), refrigerated vapor trap (Savant RVT5105); Thermo Fisher Scientific, Dreieich, Germany) and resuspended in a defined volume of 500 µl. The lipophilic fractions were resuspended in isopropanol/chloroform (3:1 v/v) containing 0.1% acetic acid, and the hydrophilic fractions were resuspended in methanol/water (50:50 v/v) containing 0.1% acetic acid. The samples were stored at -80°C until measurement. Before measurement, the samples were diluted 1:300 with the respective eluent. Each of the experimental replicates was extracted in triplicates.

### 2.8 DI-FT-ICR-MS measurements

A FT-ICR-MS (7 Tesla, SolariXR, Bruker, Bremen, Germany) with direct injection, linked to an HPLC 1260 Infinity (Agilent, Waldbronn, Germany), was used for the measurements. The eluent for the hydrophilic samples was water/methanol (50:50, v/v), and for the lipophilic samples, isopropanol/chloroform (3:1, v/v), respectively. The samples were ionized with an electrospray ionization source (pos./neg.). Two different methods for the detection of small (<1500 Da) and ultra-small molecules (<600 Da) were used, each optimized to the respective detection ranges. In total, a range of 58-3000 m/z was covered. The main parameters were nitrogen as dry gas with a temperature of 200°C and a flow rate of 4 liters/min; nitrogen was also used as nebulizer gas with a pressure of 1 bar; the time-of-flight section was set to 0.35/0.6 ms; the quadrupole mass was set to 100/200 m/z; the quadrupole RF was 2 MHz, and the sweep excitation power of the ICR-cell was 18%. All methods were calibrated before by using sodium trifluoroacetate.

### 2.9 Data handling and metabolite annotation

FT-ICR-MS data were processed by MetaboScape 5.0 (Bruker, Bremen, Germany). All datasets were recalibrated using in the sample matrix commonly occurring compounds (like glucose and amino acids) with a tolerance of <0.5 ppm. To reduce noise or false positive results, features of interest needed to be detected in at least 75% of the samples within a sample group (75% of the nine samples, i.e., 3 experimental replicates in triplicate extraction replicates of one batch at one time). If that was not the case, features were ignored and not further processed. Moreover, an intensity threshold was set to at least 10^6^.

An annotation list was created for metabolite annotation based on microbial degradation products of the plant compounds genistein (Chang & Nair, 1995; Coldham et al., 2002; Doerge et al., 2016; Gaya et al., 2016; Lee et al., 2017; Makarewicz et al., 2021; Matthies et al., 2008; Mosele et al., 2015; Schoefer et al., 2002; Soukup et al., 2023; Thompson, 2010; Wang et al., 2004), rutin, quercetin, and quercetin derivatives (Almeida et al., 2018; Aura et al., 2002; Braune et al., 2001; Di Pede et al., 2020; Duda-Chodak et al., 2015; Huang et al., 2022; Jaganath et al., 2009; Mansoorian et al., 2019; Mosele et al., 2015; Najmanová et al., 2016; Olthof et al., 2003; Parkar et al., 2013; Riva et al., 2020; Schneider et al., 1999; Schoefer et al., 2003; Serra et al., 2012; Wang et al., 2020) described in the literature. Furthermore, potential metabolites have been calculated based on Cao et al. (2015) and Traquete et al. (2022).

Features were annotated if the mass error was below 2 ppm and if the mSigma (isotope error) was below 500. For the annotation, the datasets were merged for positive and negative ion modes for each of the two mass ranges. As potential adducts, [M+H]^+^, [M+Na]^+^, and [M+K]^+^ in the positive mode and [M-H]^-^ and [M+Cl]^-^ in the negative mode were considered. The data tables were exported into R for further pre-processing. Sample replicates were combined by taking the median intensity. If a metabolite was detected by both methods (2 different mass ranges), the measurement with either the lowest number of non-detects (NA) or the highest median intensity across all samples was selected. Subsequently, the intensities of metabolites detected in the hydrophilic and lipophilic (phases) datasets were summed. NA values were replaced by zero or by the mean value if it was detected in more than 75% of the replicates.

### 2.10 Statistics metabolomics analyses with MetaboAnalyst

For the data visualization, the online tool MetaboAnalyst Version 5.0 was used (Xia et al., 2009). Modul statistical analysis (one factor), based on peak intensities, unpaired, without normalization, transformation, and scaling, was selected. In the heatmaps, the Euclidean distance method with Ward clustering algorithm was used.

## 3 RESULTS AND DISCUSSION

### 3.1 16S rRNA gene amplicon sequencing

#### 3.1.1 Changes in the bacterial communities at the level of phyla and of alpha- and beta-diversity

Both time- and treatment-dependent changes were observed. As shown in **Figure 1A**, the phyla *Firmicutes*, *Bacteroidota*, *Actinobacteriota*, *Proteobacteria*, and *Verrucomicrobiota* constituted the five main phyla in both pools. These phyla are well known to be the major phyla associated with the human gut microbiome (Rinninella et al., 2019). Already at the phylum level, it was observed that *Firmicutes*, *Bacteroidota*, and *Verrucomicrobiota* generally decreased in abundance over time, while *Actinobacteriota* and *Proteobacteria* increased during the experiment. This was probably due to the selective effect of the bacteriological growth medium used for culturing in the experiment. Furthermore, the occurrence of *Proteobacteria* seemed to be specifically supported by the plant compounds, with a minimal higher trend for rutin.

**Fig. 1.**
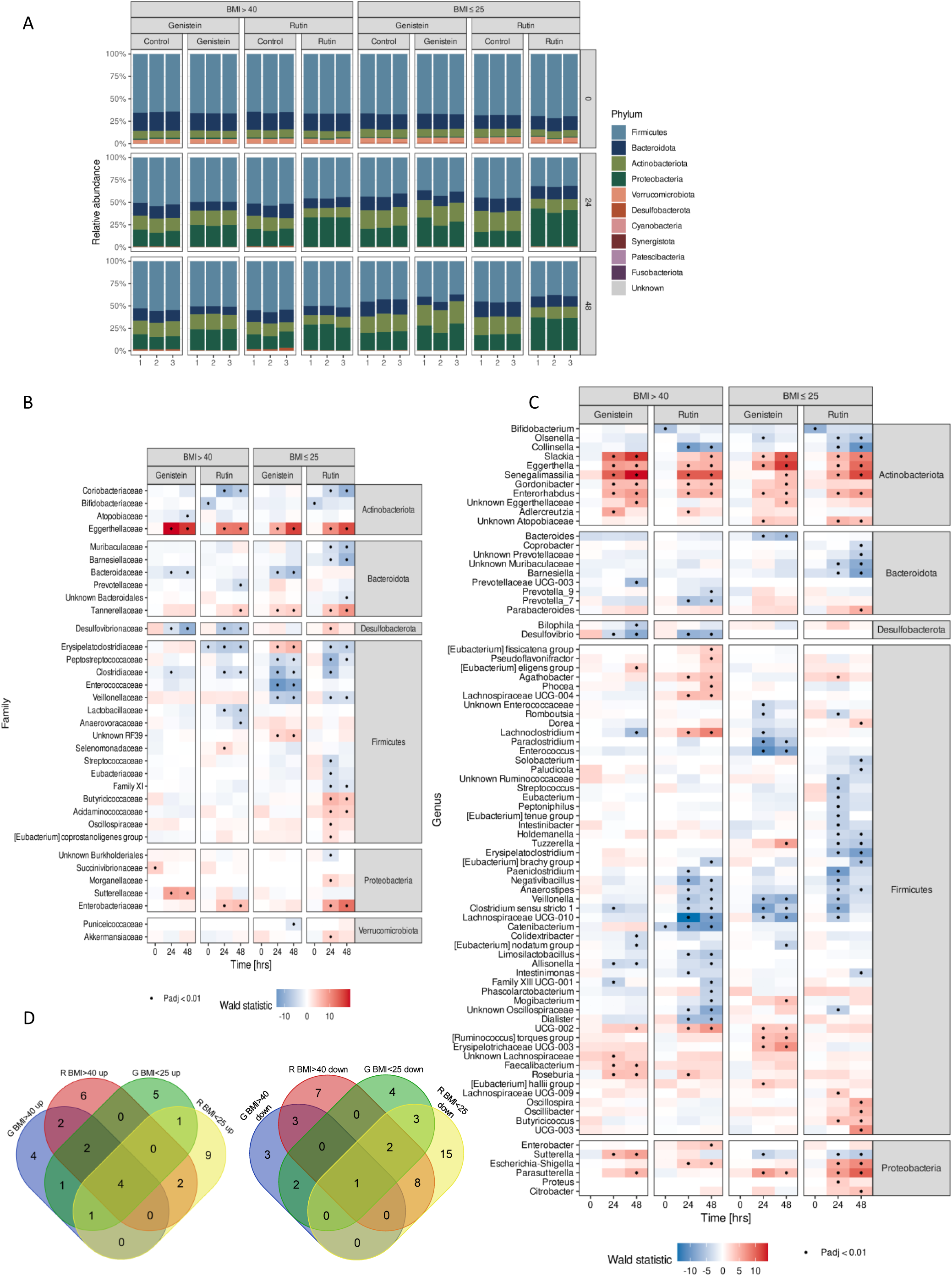
Treatment- and time-dependent changes in the bacterial communities of the two different BMI pools during the ex-vivo fecal cultivation experiments with rutin (R) and genistein (G) **A** Changes at the level of phyla, **B** changes at the level of families, **C** changes at the level of genera, and D Venn diagrams based on datasets of genera. Heatmaps show changes (red: increase and blue: decrease) in the abundance of bacteria in comparison to the corresponding controls without phytochemicals; the statistic is based on Wald, the data were p-adjusted, and families and genera were grouped by phylum.

In general, the samples of the BMI >40 pool tended to have a lower overall alpha diversity than the BMI <25 pool at 0 h. Within the first 24 h, the alpha diversity decreased but then increased again afterward in both samples treated with plant compounds and their respective controls. This was observed using 6 different algorithms (Evenness, Fisher’s alpha, Inv. Simpson div., Simpson div., Shannon div., and Species richness). At 48 h, the Shannon and Evenness diversity indices showed a significantly lower diversity in rutin-treated samples when compared to the controls in the BMI <25 pool. In addition, the Simpson div. and inverse Simpson div. indices showed significant differences in the BMI >40 pool at 24 and 48 h and in the BMI <25 pool at 24 h for both treated samples and controls **(Fig. 2A-F)**. Beta diversity was also different between the two BMI pools. Compared to the start of cultivation (0 h), the differences in beta diversity between the 24- and 48-hour samples were more distinct, as shown by a clear separate clustering **(Fig. 2G)**; this trend applied equally to both phytochemicals, again indicating a potential bacteriological growth medium effect. The genistein-treated samples differed from the controls at 24 and 48 h, whereas the rutin-treated samples differed in the BMI <25 pool at the same periods.

**Fig. 2.**
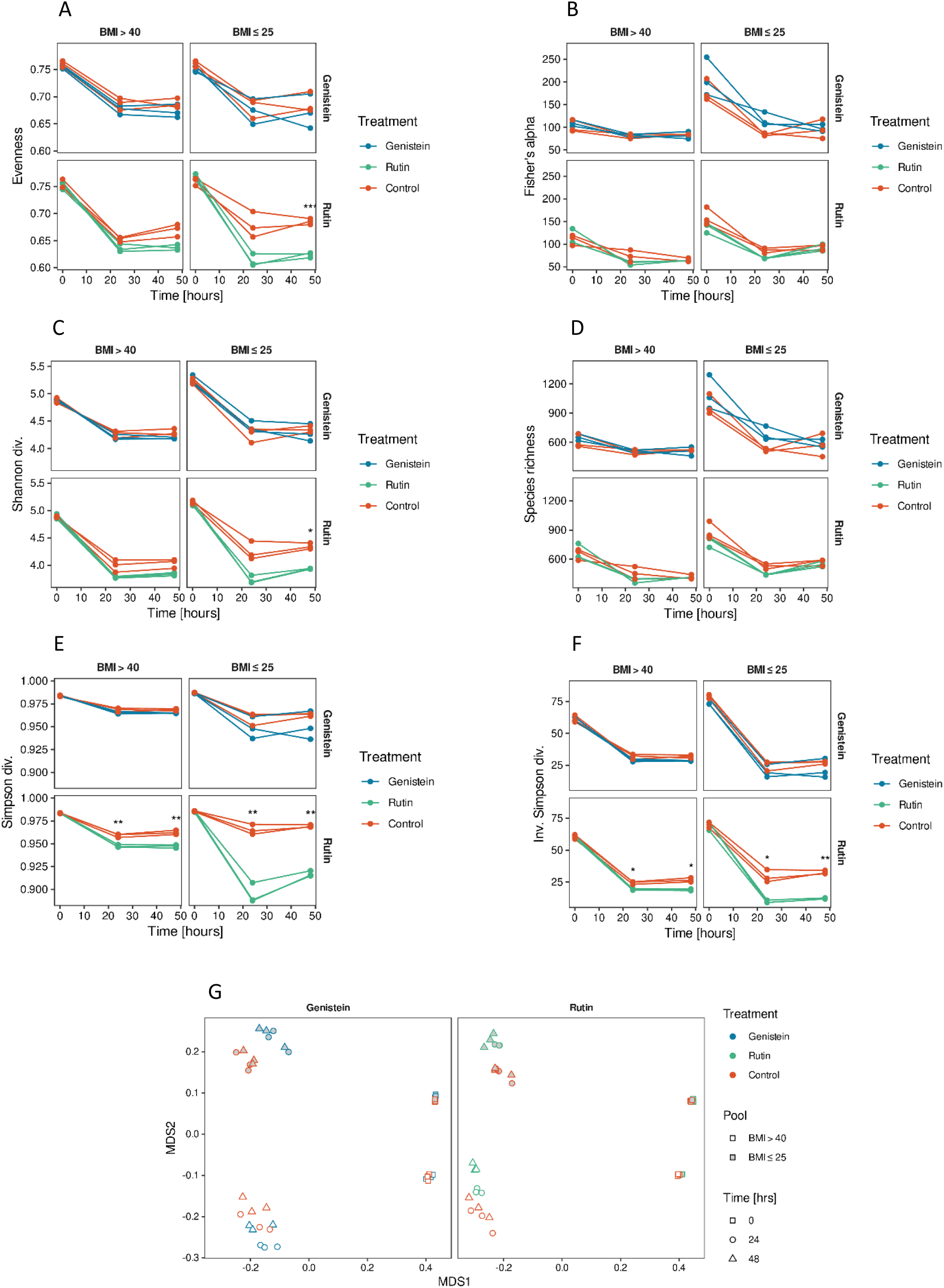
Treatment and time-dependent changes in α-/β-diversity of the bacterial communities. **A-F** changes at the level of α-diversity over time as six different diversity indexes, separated by the two different pools, BMI >40 and BMI ≤25, and the two different treatments, rutin and genistein. One graph corresponds to one experimental replicate. **G** time and treatment-dependent changes in β-diversity. In all figures **(A-G)**, genistein-treated samples are shown in blue, rutin-treated samples in green, and the corresponding controls in orange.

#### 3.1.2 Changes in the bacterial communities at family and genus levels

A total of 34 families and 83 genera were significantly affected (Wald p <0.01) by the rutin and genistein treatments **(Fig. 1B/C).** Of these, 13 families were significantly stimulated, while 19 were significantly inhibited by either one or both of the compounds. For two families, an opposite effect was observed, as these were either up- or down-regulated by one phytochemical while the opposite occurred with the other **(Fig. 1B)**.

The 16S rRNA gene metagenomics data revealed effects on family abundance caused equally by both plant compounds in both BMI pools. The most prominent and obvious effect was that both rutin and genistein caused a significant decrease in the abundance of *Clostridiaceae* and a significant increase in the abundance of *Eggerthellaceae* in both BMI pools **(Fig. 1A)**.

Accordingly, at the genus level, four genera belonging to the *Eggerthellaceae* were stimulated to increase in abundance by both plant compounds in both of the BMI pools, i.e., *Enterorhabdus*, *Eggerthella*, *Slackia*, and *Senegalimassilia*, while one was decreased (*Clostridium sensu stricto 1*). The genus *Enterorhabdus* is a heterotypic synonym of *Adlercreutzia* (Nouioui et al., 2018). Since these genera can be considered to be taxonomically the same, these bacteria were also stimulated by both plant compounds in both BMI groups **(Fig. 1B)**. In addition, the genus *Gordonibacter* was also stimulated by both rutin and genistein. However, only genistein significantly stimulated *Gordonibacter* in both BMI groups, whereas rutin showed a significant effect only in the BMI >40 group, with a positive trend observed in the BMI <25 group as well.

A systematic review of the composition of the gut microbiome in obese and non-obese persons found two studies that reporting bacteria of the genus *Eggerthella* were less abundant in obese persons (Pinart et al., 2021), our studies indicated that plant compounds increased their abundance in feces of both BMI groups. This increase may be explained by the fact that certain species of these genera/families are capable of metabolizing of plant polyphenols and thus may be nutritionally selected for. Specific bacteria belonging to the *Eggerthellaceae* are well known to be involved in the metabolism of plant compounds, such as the isoflavone genistein used in this study (Braune & Blaut, 2016; Makarewicz et al., 2021; Setchell & Clerici, 2010; Soukup et al., 2023), which may explain their selective effect. Rutin (quercetin-3-O-rutinoside) is a flavonol glycoside composed of quercetin and rutinose, a disaccharide of rhamnose and glucose. The *Eggerthellaceae* have not been reported to possess an α-rhamnosidase enzyme that removes the sugar moiety from the quercetin aglycone. A search for this gene in the NCBI gene database (Sayers et al., 2022) did not show the presence of this enzyme in the genomic sequences of genera belonging to this family (results not shown). However, it is known that bacteria belonging to the genus *Eggerthella* can metabolize quercetin (Braune & Blaut, 2016), and this may explain the increase in abundance of these bacteria also in the samples treated with rutin. This bacterial genus possesses specific enzymes capable of breaking down the structure of quercetin, which may be the result of co-evolution with the host. *Eggerthella* is part of the human gut microbiota, which interacts closely with the host’s diet and immune system. The ability to degrade quercetin could, therefore, represent an evolutionary adaptation that benefits both the bacteria and the host. Given the stimulatory effect of quercetin on *Eggerthella* growth, it is likely that the polyphenol serves as a carbon and energy source for the bacteria. The subsequent metabolites released during the degradation process may influence the composition of the microbiota, potentially modulating the competitive and synergetic bacteria present. The antimicrobial properties of polyphenols suggest a possible detoxification role for the degradation process. However, given the growth stimulation of *Eggerthella* by quercetin, it is reasonable to assume that its utilization as an energy source is the most likely explanation (Rodríguez-Daza et al., 2021). The stimulation of other bacteria could therefore also be linked to cross-feeding interactions.

The increase in abundance of bacteria belonging to the *Eggerthellaceae* may be considered beneficial, as these bacteria are known to metabolize polyphenols such as quercetin and genistein into bioactive substances. As a systematic review found that these bacteria were reduced in the fecal microbiota of obese individuals, an intervention with plant compounds might be able to restore the abundance of *Eggerthellaceae*. However, it should be that *Eggerthella lenta* has also been described as an opportunistic pathogen involved in bacteremia (Gardiner et al., 2015; Ugarte-Torres et al., 2018), has been found to be more abundant in type 2 diabetes patients (Koh et al., 2018; Qin et al., 2012), and produces imidazole propionate, which interferes with insulin signaling (Koh et al., 2018). *Eggerthella lenta* is also capable of metabolizing the cardiac drug digoxin (Koppel et al., 2018). Thus, care should be taken to define the conditions under which such bacteria could be used in potential intervention studies.

Another general effect was that both polyphenolic plant compounds showed a clear inhibitory effect on the family *Clostridiaceae*, specifically on bacteria of the genus *Clostridium* **(Fig. 1A/B).** While the exact reason for this is not clear, it is known that tea polyphenols, for example, are inhibitory to *Clostridium* spp. (Ahn et al., 1991). Furthermore, the *Bifidobacteriaceae*, specifically the genus *Bifidobacterium*, and *Coriobacteriaceae*, specifically the genus *Collinsella*, were significantly inhibited by rutin treatment in both BMI pools. On the other hand, rutin also positively affected the abundance of *Enterobacteriaceae* in both BMI pools. At the genus level, this pertained to the genus *Escherichia-Shigella* in both BMI pools, as well as *Enterobacter* in the BMI >40 pool and *Citrobacter* in the BMI >25 pool **(Fig. 1B)**. Bifidobacteria have previously been shown to be unable to hydrolyze rutin (Amaretti et al., 2015), which may indicate a selective disadvantage for these bacteria. A study by Gwiazdowska et al. (2015)showed that while rutin was shown to be stimulate bifidobacterial growth, the quercetin aglycone inhibited the growth of certain species such as *B. bifidum*, which may explain the inhibition of bifidobacteria in the fecal samples treated with rutin in this study. This may be considered to be a detrimental effect, as bifidobacteria are well known for their role as probiotics, with beneficial effects such as anti-cancer, anti-inflammatory, anti-infective, and immune regulatory effects (Chen et al., 2021). The increase in *Enterobacteriaceae* in the presence of rutin can also be considered as detrimental, as enterobacterial lipopolysaccharide (LPS), which is an important component of the bacterial cell wall, plays a crucial role in the progression of inflammation and insulin resistance (Barlow et al., 2015). Enteric LPS, also considered an endotoxin, can enter the bloodstream through the damaged intestinal mucosa and then cause systemic inflammation (Xu et al., 2018). In addition, some of the enterobacteria are well known to be opportunistic pathogens.

Also, of note was a decrease in the abundance of *Lactobacillaceae*, specifically the genus *Limosilactobacillus,* in rutin-treated fecal samples of the BMI >40 group and a decrease in the abundance of *Enterococcaceae* (specifically the genus *Enterococcus*) in genistein-treated fecal samples of the BMI <25 group **(Fig. 1A/B)**. While both limosilactobacilli (e.g., *L. reuteri*, *L. mucosae*) and enterococci (e.g., *E. faecium, E. faecalis*) are lactic acid bacteria and typical for the human gastrointestinal tract, positive associations are usually attributed to lactobacilli rather than the enterococci. This is because the former are well known for their probiotic properties, while the latter include pathogenic strains that harbor virulence factors, toxins, or antibiotic resistance genes (Colautti et al., 2022). The genus *L. reuteri* is currently used as a probiotic as it has been associated with gut health due to its anti-inflammatory effects, regulation of the gut microbiota, and improvement of the gut barrier (Yu et al., 2023). Genistein also increased the abundance of bacteria belonging to the genera *Roseburia* and *Faecalibacterium* in the BMI >40 group. Like lactobacilli, *Roseburia* spp. and *Faecalibacterium prausnitzii* are generally considered beneficial bacteria that are involved in anti-inflammatory and antioxidant processes (Man et al., 2020). However, *Roseburia* is considered to be dysregulated in the context of obesity (Pinart et al., 2021), so the growth-stimulating effect of both plant compounds in the BMI >40 pool is rather negative.

Rutin was previously found to enrich *Lachnospiraceae* (*Lachnoclostridium* and *Eisenbergiella*), *Enterobacteriaceae*, *Tannerellaceae*, and *Erysipelotrichaceae* species in the human fecal microbiota, and it was observed that *Enterobacteriaceae* were associated with the conversion of rutin to quercetin-3-glucoside and *Lachnospiraceae* were associated with the conversion to quercetin (Riva et al., 2020). An increase in *Enterobacteriaceae*, *Tannerellaceae, and* some genera of *Lachnospiraceae* in rutin-treated fecal microbiota was also observed in our study for both BMI groups, confirming the results of Riva et al. (2020) **(Fig. 1A)**. At the genus level, an increase of bacteria belonging to the *Tannerellaceae* genus *Parabacteroides* was observed, at least for the rutin-treated samples of the BMI <25 group. Increases in several genera of *Lachnospiraceae*, including the genus *Agathobacter* in both BMI groups, *Roseburia* in the BMI >40 group, *Lachnoclostridium* in the BMI >40 group, and *Dorea* in the BMI >25 groups, also occurred as a result of the presence of rutin in the fecal samples **(Fig. 1B)**. Interestingly, the abundance of bacteria belonging to the genus *Sutterella* decreased in the fecal samples of the BMI <25 groups to which either of the plant compounds was added. Conversely, the abundance of bacteria belonging to the genus *Sutterella* increased in the genistein-treated fecal samples from the BMI >40 group. A systematic review showed that *Sutterella* was significantly more abundant in obese individuals than in non-obese individuals (Pinart et al., 2021). Therefore, in this BMI group, genistein led to an even greater increase in the abundance of these bacteria. In our study, *Parasutterella* abundance was also increased by genistein in fecal samples from both BMI groups, and the rutin-treated BMI <25 group samples **(Fig. 1B)**. *Parasutterella* was previously identified to be associated with both BMI and type 2 diabetes (Henneke et al., 2022).

Other effects of rutin or genistein on changes at either the family or genus level occurred that were either BMI group or specific plant compound treatment-related effects. For example, bacteria belonging to the family *Bacteroidaceae* were significantly inhibited by genistein in both of the BMI pools. Both rutin and genistein induced a response only in the BMI <25 pools, resulting in a decrease of *Peptostreptococcaceae* and *Veillonellaceae*. Rutin and genistein also increased the *Tannerellaceae* in the BMI <25 pools, and bacteria belonging to this family increased in abundance in the BMI >40 pool with rutin. Moreover, very specific effects were observed, with only one BMI pool showing a significant specific response to only one of the plant compounds **(see Fig. 1A/B)**. A particularly beneficial aspect is the inhibitory effect of rutin on the genera *Prevotella* 7 and 9, *Cantenibacterium*, and *Dialister* within the BMI >40 pools due to their dysregulation in obesity.

The detrimental effects, such as those observed for bifidobacteria and lactobacilli, may be countered by employing different approaches. In individuals with a healthy microbial composition, this issue could potentially be mitigated by administering dietary fiber, which promotes the growth of these beneficial bacteria (“bifidogenic effect”). Alternatively, if microbial metabolites are identified as the source of positive effects, directly utilizing or supplementing these metabolites could serve as a more targeted strategy. A combination of polyphenols could also be used, for example, including one known to have a growth-stimulating effect on the bacterial genera that are reduced in our study. Additionally, probiotics with lactobacilli and bifidobacteria could be supplemented or obtained through fermented foods.

### 3.2 Metabolomics analyses

#### 3.2.1 Degradation and metabolization of rutin

In a semi-targeted analysis, metabolite profiles were screened for previously described degradation products (**see. 2.9**) as well as for not-yet-described transformation products of rutin that would result from methylations, sulfations, and dihydroxylations. The following numerical data were calculated at the molecular formula level to avoid overrepresentation of isobaric compounds.

A total of 46 degradation and transformation products associated with a clear rutin-dependent effect were tentatively identified **(Fig. 3)**. Of the 46, 25 showed a treatment-dependent trend in both pools, i.e., they had a higher intensity in the treatments (samples containing rutin) than in the corresponding controls or occurred only in the treatments. To our knowledge, this is the first time that such a high number of degradation and transformation products of microbial origin has been detected in a single experiment.

**Fig. 3.**
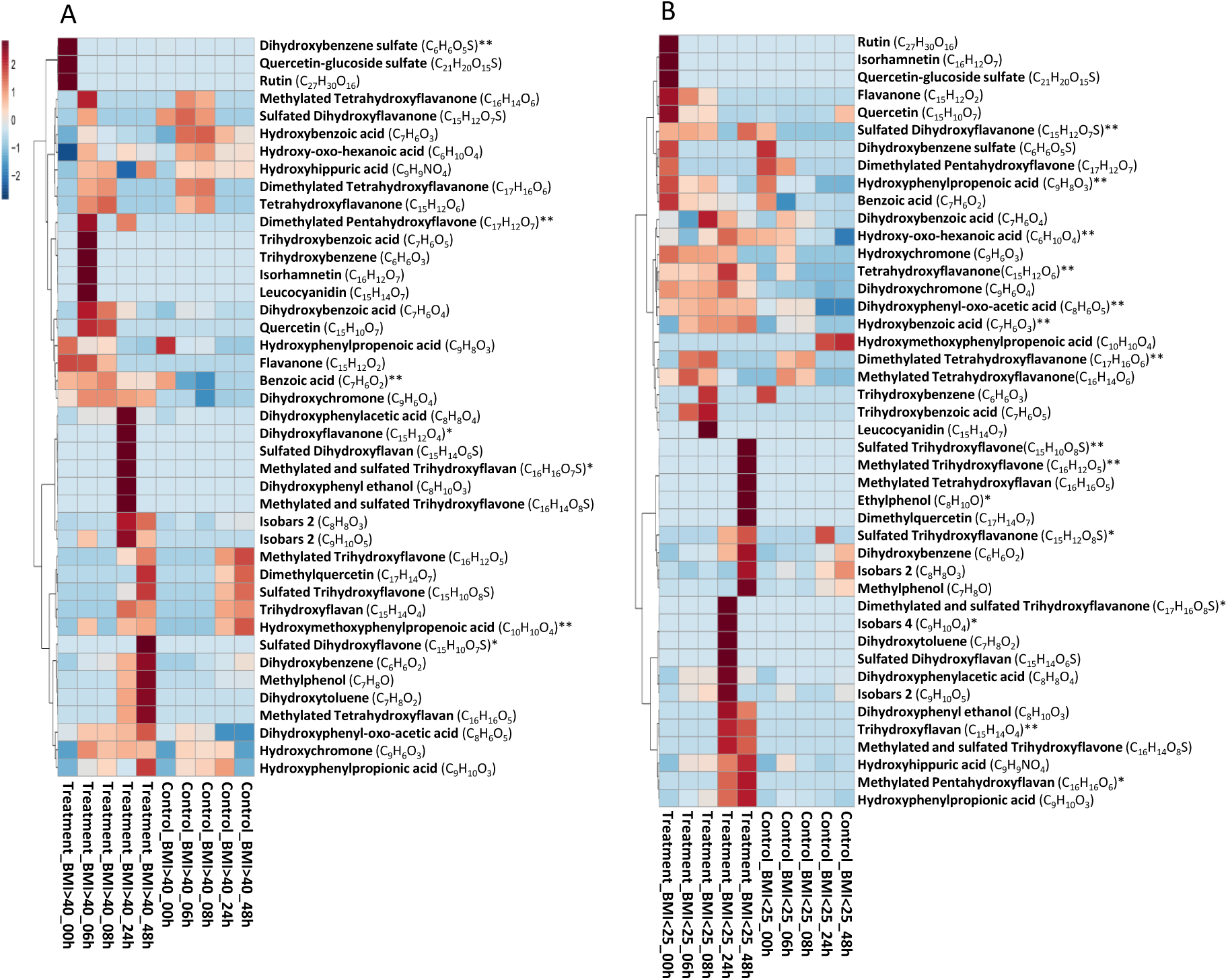
Degradation and transformation of rutin. Heatmap clustering analysis performed with MetaboAnalyst 5.0 using the obtained merged MS data. Calculation based on mean values of the experimental replicates. Analysis was performed using the Euclidean distance method with the Ward clustering algorithm. **A** treatment vs. control pool BMI >40, **B** treatment vs. control pool BMI <25 for the respective times of 0, 6, 8, 24, 48h. Reported are only metabolites with a treatment-dependent trend in at least one of the two pools in comparison to the control. ** trend in the corresponding treatment only; * occurrence of the compound in the corresponding treatment only. Isobaric compounds **C_8_H_8_O_3_** hydroxyphenylacetic acid or hydroxymethoxybenzoic acid; isobaric compounds **C_9_H_10_O_5_** hydroxydimethoxybezoic acid or methoxyhydroxyphenylhydroxyacetic acid; **C_9_H_10_O_4_** hydroxyphenylhydroxypropionic acid, methoxyhydroxyphenylacetic acid, dihydroxyphenylpropionic acid or dimethoxybenzoic acid.

In both BMI pools, the precursor rutin was degraded within 6 hours. In the first step, the sugar residue was split off. From the presence of the tentatively identified metabolites quercetin and quercetin-glucoside-sulfate, it could be deduced that both the complete sugar residue and initially only a part of the sugar residue were split off. The entire rutinoside could be cleaved through the enzyme ß-rutinosidase or only partially by first an α-mannosidase and then by a ß-glucosidase. However, the second degradation pathway is more common, as only a few bacteria possess a ß-rutinosidase (Riva et al., 2020). The degradation of rutin to quercetin, or of rutin via quercetin glucosides to quercetin, could also be shown by Riva et al. (2020).

Starting with the aglycone quercetin, various other degradation and transformation products were observed in the subsequent course of the experiment. Numerous benzoic acid compounds were detected, which result from the destabilization and cleavage of the C-ring and are products of the A- and B-ring. These have also been described previously (Almeida et al., 2018; Aura et al., 2002; A. Braune et al., 2001; Di Pede et al., 2020; Duda-Chodak et al., 2015; Jaganath et al., 2009; Mansoorian et al., 2019; Mosele et al., 2015; Najmanová et al., 2016; Olthof et al., 2003; Parkar et al., 2013; Schneider et al., 1999; Schoefer et al., 2003; Serra et al., 2012; Wang et al., 2020) and include, for example, the benzoic acids trihydroxybenzoic acid, dihydroxybenzoic acid, hydroxybenzoic acid, and benzoic acid, as well as dihydroxyphenylacetic acid and hydroxyphenylacetic acid, which are formed by simple dehydroxylation.

In addition to these A- and B-ring products, compounds were also found in both BMI pools in which no C-ring cleavage occurred and the A-, B- and C-rings remained intact, and in which the A- and C-ring remained intact and the B-ring was cleaved off. Products with an intact A-, B-, and C-ring included isorhamnetin (methylquercetin), dimethylquercetin, leucocyanidin, and flavanone, as well as several previously undescribed compounds. Compounds with an intact A- and C-ring were dihydroxychromones and hydroxychromones; the latter was also described by Huang et al. (2022) for the degradation of rutin by *B. uniformis*.

As reviewed by Zhao et al. (2022), bacteria have a broad spectrum of enzymes, in addition to glucosidases, hydrolases, oxidoreductases, lyases, and transferases. In particular, Cao et al. (2015) described hydroxylation and dehydroxylation, O-methylation and O-demethylation, glycosylation and deglycosylation, hydrogenation and dehydrogenation, as well as sulfation in the context of microbial biotransformation of flavonoids. Taking these possible reactions into account, molecular formulae and mass differences were calculated according to the approach of Traquete et al. (2022) for the prediction of possible transformation products, which were then specifically searched for. It was also taken into account that multiple reactions may occur, so that a molecule may, e.g., be both methylated and sulfated, or dehydroxylated or methylated multiple times. In this context, in addition to the above-mentioned metabolites flavanone and dihydroxychromone, which in our opinion have not yet been described as degradation products of rutin, another 17 undescribed transformation products were tentatively identified.

Starting from quercetin (C_15_H_10_O_7_), a metabolite with the molecular formula C_15_H_12_O_6_ was detected. This may have been formed by hydrogenation of quercetin to dehydroquercetin (C_15_H_12_O_7_; not detected) and subsequent additional dehydroxylation. This may represent the tetrahydroxyflavanone eriodictyol. Starting from eriodictyol, the metabolite with the molecular formula C_15_H_12_O_4_ could also be formed by double dehydroxylation; this could be the dihydroxyflavanone dihydrochrysin. Neither eriodyctol nor dihydrochrysin have been described in connection with rutin. From dihydrochrysin, dihydrochrysin sulfate (C_15_H_12_O_7_S) may have been formed by sulfation. The two unknown metabolites with the molecular formulae C_16_H_14_O_6_ and C_17_H_16_O_6_ could have been formed from eriodictyol by methylation and dimethylation.

Quercetin itself is a pentahydroxyflavone, the detected leucocyanidin is a hexahydroxyflavan, and the previously mentioned dihydroquercetin is a pentahydroxyflavanone.

Based on possible dehydroxylations, methylations, and sulfations, the remaining metabolites could analogously be a sulfated trihydroxyflavone (C_15_H_10_O_8_S) and a sulfated dihydroxyflavone (C_15_H_10_O_7_S), furthermore, a dimethylated pentahydroxyflavone (C_17_H_12_O_7_) and a methylated trihydroxyflavone (C_16_H_12_O_5_). In addition, a sulfated trihydroxyflavanone (C_15_H_12_O_8_S) and a sulfated and methylated trihydroxyflavone (C_16_H_14_O_8_S) were tentatively identified. Additionally, a trihydroxyflavan (C_15_H_14_O_4_), a methylated pentahydroxyflavan (C_16_H_16_O_6_), a methylated tetrahydroxyflavan (C_16_H_16_O_5_), a sulfated dihydroxyflavan (C_15_H_14_O_6_S), a methylated and sulfated trihydroxyflavan (C_16_H_16_O_7_S), and a sulfated and dimethylated trihydroxyflavanone (C_17_H_16_O_8_S) were found.

Dihydroxyflavanone, sulfated dihydroxyflavone, and methylated and sulfated trihydroxyflavan were only found in samples from the BMI >40 pool. Ethylphenol, pentahydroxyflavan, the dimethylated and sulfated trihydroxyflavanone, and the sulfated trihydroxyflavanone, as well as a metabolite with the molecular formula C_9_H_10_O_4_, corresponding to the 4 different isobaric compounds hydroxyphenylhydroxypropionic acid, methoxyhydroxyphenylacetic acid, dihydroxyphenylpropionic acid, or dimethoxybenzoic acid, could only be detected in the BMI <25 pools. In principle, it is assumed that although microbiomes may differ in their composition, the core functions remain approximately the same, but individual metabolism or differences in metabolic profiles may occur. Different bacterial metabolic profiles or, metabotypes are known, with regard to the phenolic metabolism: fast, medium, and slow converters; former and non-former; and low and high producers (Hu et al., 2024). Metabolomic analyses are snapshots and represent the current situation at the time of sampling, so it is also be possible that certain metabolites were not present at the time of sampling. The fact that the samples differ in their rate of metabolism is also evident from different metabolites. For example, leucocyanidine is formed earlier in the BMI >40 samples than in the BMI <25 samples, while dihydroxytoluene is formed earlier in the BMI <25 samples than in the BMI >40 samples.

#### 3.2.2 Degradation and metabolization of genistein

For genistein, a total of 23 degradation and transformation products were tentatively identified **(Fig. 4)**. This wide range has, to our knowledge, not been described before. 13 of the 23 compounds were found in both pools, BMI >40 and BMI <25 pools, while another 3 were found only in the BMI >40 pool and 5 compounds only in the BMI <25 pool. Dihydrogenistein (C_15_H_12_O_5_) was only found in the BMI <25, while ethylphenol (C8H10O) was found only in the BMI >40 pool. Similar to rutin, genistein was metabolized within 6 hours in both BMI pools.

**Fig. 4.**
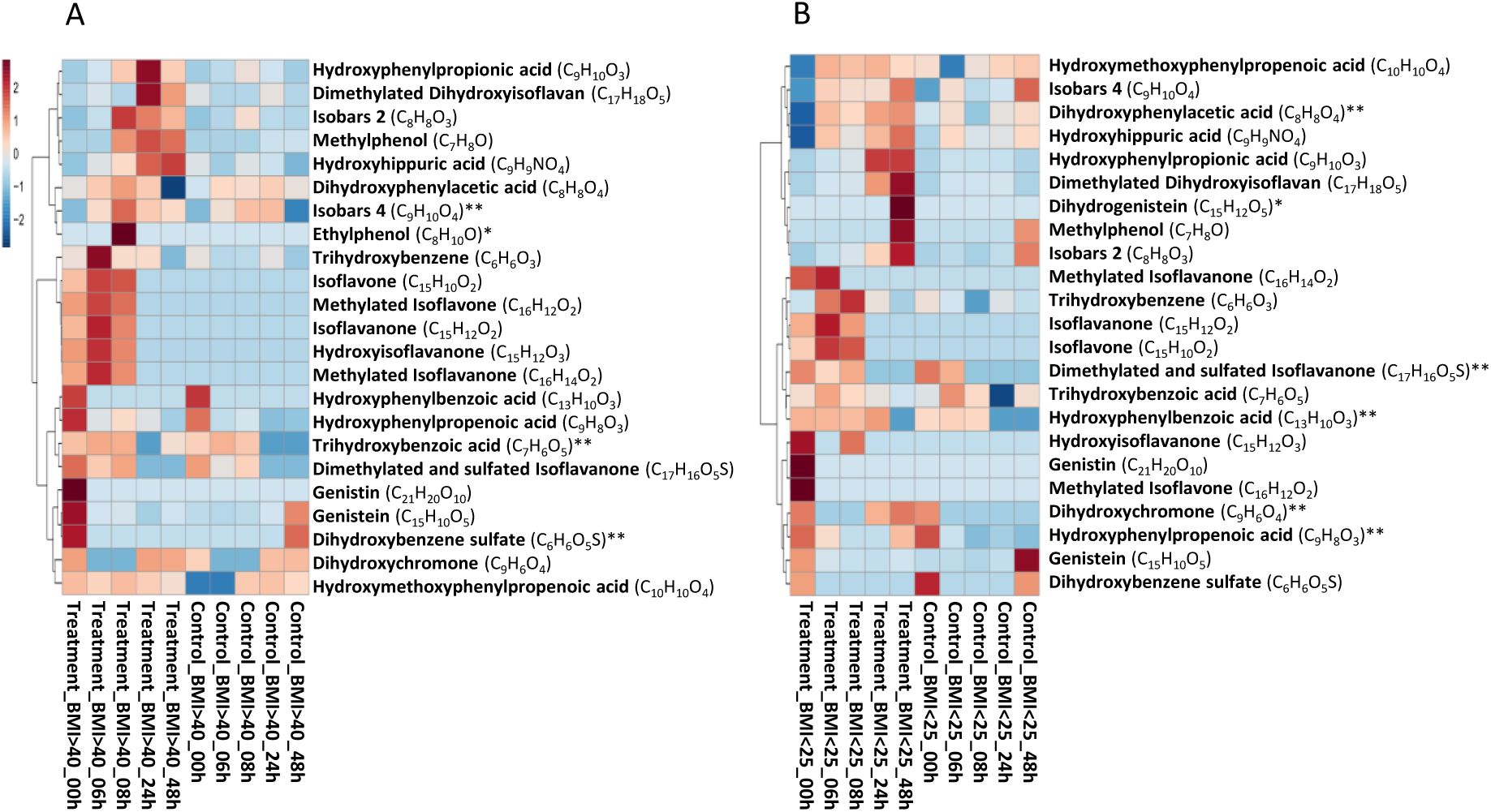
Degradation and transformation of genistein. Heatmap clustering analysis was performed with MetaboAnalyst 5.0 using the obtained merged MS data. Calculation based on mean values of the experimental replicates. Analysis was performed using the Euclidean distance method with the Ward clustering algorithm. **A** treatment vs. control pool BMI >40, **B** treatment vs. control pool BMI <25 for the respective times of 0, 6, 8, 24, 48h. Reported are only metabolites with a treatment-dependent trend in at least one of the two pools in comparison to the control. ** trend in the corresponding treatment only; * occurrence of the compound in the corresponding treatment only. Isobaric compounds **C_8_H_8_O_3_** hydroxyphenylacetic acid or hydroxymethoxybenzoic acid; **C_9_H_10_O_4_** hydroxyphenylhydroxypropionic acid, methoxyhydroxyphenylacetic acid, dihydroxyphenylpropionic acid, or dimethoxybenzoic acid.

As shown for rutin, metabolites were found that either still had the intact basic structure of the flavone consisting of A-, B- and C-ring (e.g., dihydrogenistein), or of A- and C-ring (e.g., dihydroxychromone), or resulted from the C-ring cleavage and carried elements of the A- or B-ring. These include, for example, methylphenol and ethylphenol, hydroxyphenylpropionic acid, dihydroxyphenylacetic acid, and trihydroxybenzene, which have also been previously described by Coldham et al., 2002; Doerge et al., 2016; Gaya et al., 2016; Lee et al., 2017; Makarewicz et al., 2021; Mosele et al., 2015; Schoefer et al., 2002; Soukup et al., 2023; Thompson, 2010; Wang et al., 2004. Dihydroxychromone has not been previously detected as a degradation product of genistein. The same applies to the tentatively identified isoflavone. Another interesting aspect is the very late appearance of dihydrogenistein at 48h only in the BMI <25 pools. This suggests that genistein is not only degraded via dihydrogenistein and then with subsequent C-ring cleavage, but that further degradation pathways must be involved since characteristic A-ring and B-ring metabolites are already formed beforehand. In analogy to rutin, novel transformation products were found in this study that have not been described before. These include a metabolite with the molecular formula C_17_H_18_O_5_, which is proposed to be a dimethylated dihydroxyisoflavan. The metabolites with the molecular formulae C_15_H_12_O_3_ and C_15_H_12_O_2_ could resulted from dehydrogenistein by multiple dehydroxylation. The metabolites with the molecular formulae C_16_H_14_O_2_ and C_17_H_16_O_5_S could drived from the metabolite with the molecular formula C_15_H_12_O_2_ after subsequent methylation (C_16_H_14_O_2_) and dimethylation and sulfation (C_17_H_16_O_5_S). The metabolite with the molecular formula C_16_H_12_O_2_ may be result of multiple dehydroxylation of genistein followed by methylation.

For both phytochemical treatments, rutin and genistein, the first degradation and transformation products could be detected already at 0 h; this could be due to the sampling, as the sampling was not performed directly, but within the first 15 min. As described by Walle et al. (2005) and Kahle et al. (2011), the first degradation products can already be formed after 5 min in an *in vitro* system for saliva.

It is also interesting to note that in addition to genistein, genistin was also detected in the 0 h samples. Genistin is the glycosylated form of genistein. It could not be detected in the stock solution, indicating that the glycosylation must be of microbial origin. Bacteria can transfer sugar residues from UDP-glucose using glycosyltransferases, this was also observed by Rabausch et al. (2013) for various polyphenols and described as a measure of detoxification, analogous to hydroxylation, sulfation, and methylation of phase I+II of biotransformation, which pursue the same goal. Microbial biotransformation thus opens up a wide field of potential new bioactive compounds.

## 4 CONCLUSION

Polyphenols affect the human gut microbiome at the level of composition and diversity, resulting in wide variation in degradation spectrum of plant compounds. Different polyphenol-related effects were observed, including polyphenol-specific effects, i.e., effects caused by both polyphenols in both BMI pools, pool-specific effects, i.e., effects caused by both plant substances within one BMI pool, genistein-specific effects, rutin-specific effects, and exclusively individual effects, i.e., effects caused by one plant substance specifically within one BMI pool. Furthermore, it was shown that polyphenols were able to stabilize and stimulate the growth of bacteria considered to be dysregulated in the context of obesity. In addition, numerous microbial degradation and transformation products of polyphenols, were tentatively identified, some of which are still unknown.

A diet rich in plant compounds, especially polyphenols, may affect the microbial community and thus, potentially the host physiology. Responses to diet can be highly individual, which may lead to completely new opportunities in personalized nutrition. The new transformation products raise new health-related questions and perspectives, as the bioactivity of these metabolites is still unknown. Furthermore, the methodological approach adopted in this study can be used to evaluate other plant compounds, dietary constituents, or drugs. In particular, the effect on the microbiota can also be tested for individuals or other diseases that are associated with a dysregulated microbial composition.

## Data availability statement

The original contributions presented in the study are included in the article/ supplementary material, further inquiries can be directed to the corresponding author. Paired-end 16S rRNA gene sequencing data are available at the European Nucleotide Archive (ENA, https://www.ebi.ac.uk/ena/browser/, accession number: PRJEB76448).

## Funding

This project was funded by the Federal Ministry of Food and Agriculture and was part of the project Gut Metabotypes as Biomarkers for Nutrition and Health (BioNUGUT) Grant No. 2816ERA13E.

## Acknowledgments

We thank Lena Westphal for her support in conducting the microbiology experiments, Adrian Prager for his support in the preparation of sequencing libraries, and Meike Pfeiler for her support in sample preparation for the metabolomics analyses.

## Declaration of interest

The authors declare that the research was conducted in the absence of any commercial or financial relationships that could be construed as a potential conflict of interest.

## Declaration of generative AI in scientific writing

The authors declare that AI was not used to write any parts of the manuscript.

## Notes

### Competing Interest Statement

The authors have declared no competing interest.

