## Supplement Fig. S1 for "Degradation of rutin and genistein and the effect on human bacterial fecal populations of obese and non-obese"

A

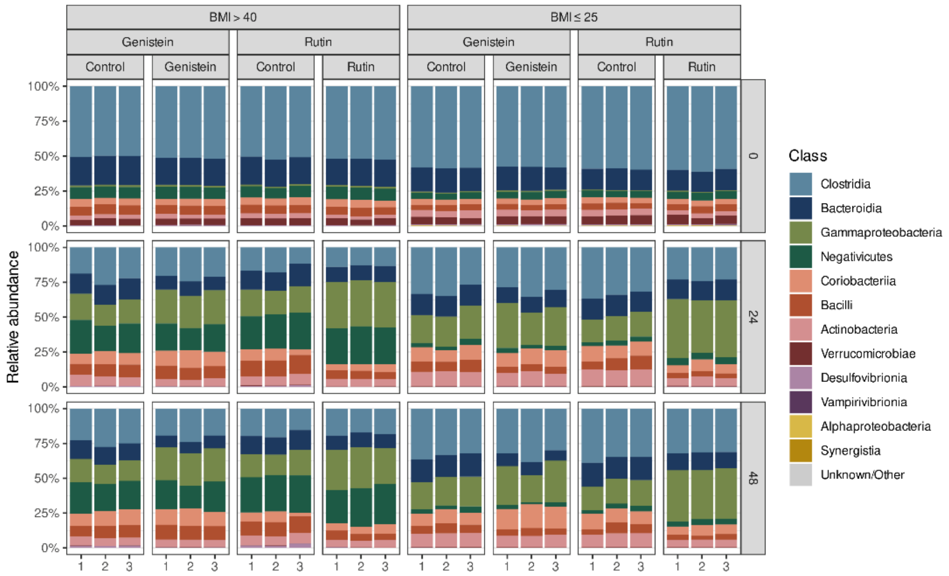
BMI > 40
BMI ≤ 25
Genistein
Rutin
Control
Genistein
Control
Rutin
100%
75%
50%
25%
0%
Relative abundance
0
24
48
1 2 3
Class
Clostridia
Bacteroidia
Gammaproteobacteria
Negativicutes
Coriobacteriia
Bacilli
Actinobacteria
Verrucomicrobiae
Desulfovibrionia
Vampirivibrionia
Alphaproteobacteria
Synergistia
Unknown/Other

B

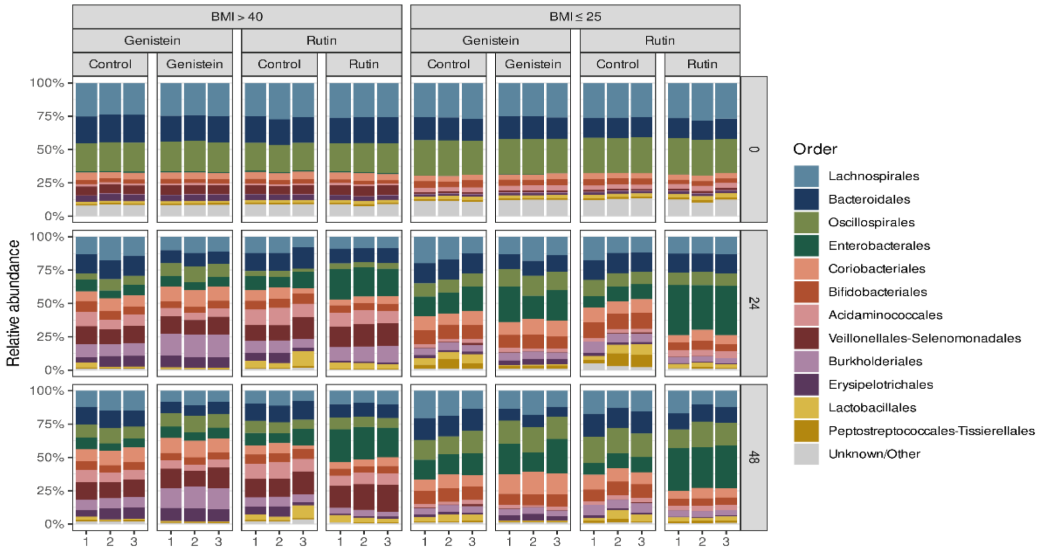
BMI > 40
BMI ≤ 25
Genistein
Rutin
Control
Genistein
Control
Rutin
100%
75%
50%
25%
0%
Relative abundance
0
24
48
1 2 3
Order
Lachnospirales
Bacteroidales
Oscillospirales
Enterobacterales
Coriobacteriales
Bifidobacteriales
Acidaminococcales
Veillonellales-Selenomonadales
Burkholderiales
Erysipelotrichales
Lactobacillales
Peptostreptococcales-Tissierellales
Unknown/Other

C

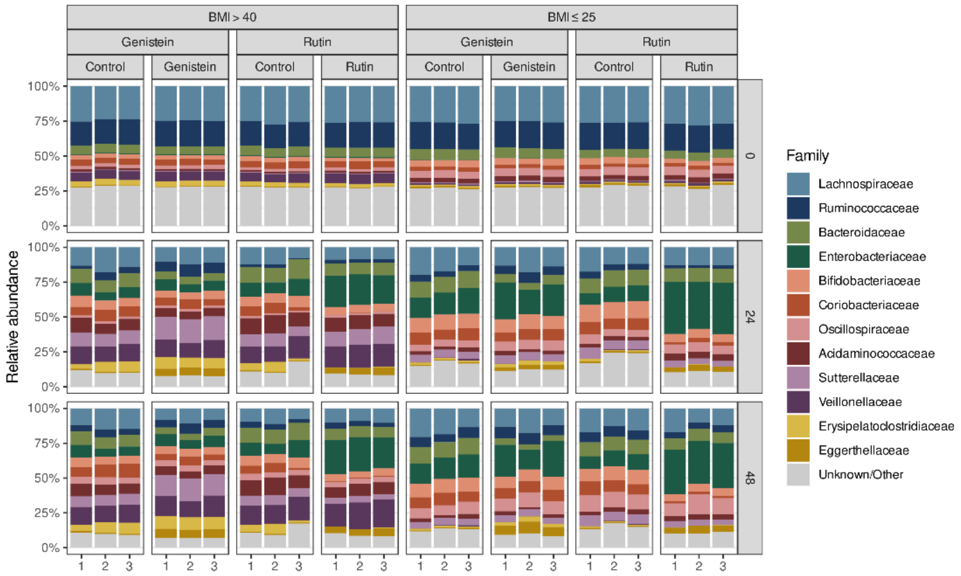
BMI > 40
BMI ≤ 25
Genistein
Rutin
Control
Genistein
Control
Rutin
100%
75%
50%
25%
0%
Relative abundance
0
24
48
1 2 3
Family
Lachnospiraceae
Ruminococcaceae
Bacteroidaceae
Enterobacteriaceae
Bifidobacteriaceae
Coriobacteriaceae
Oscillospiraceae
Acidaminococcaceae
Sutterellaceae
Veillonellaceae
Erysipelatoclostridiaceae
Eggerthellaceae
Unknown/Other

D

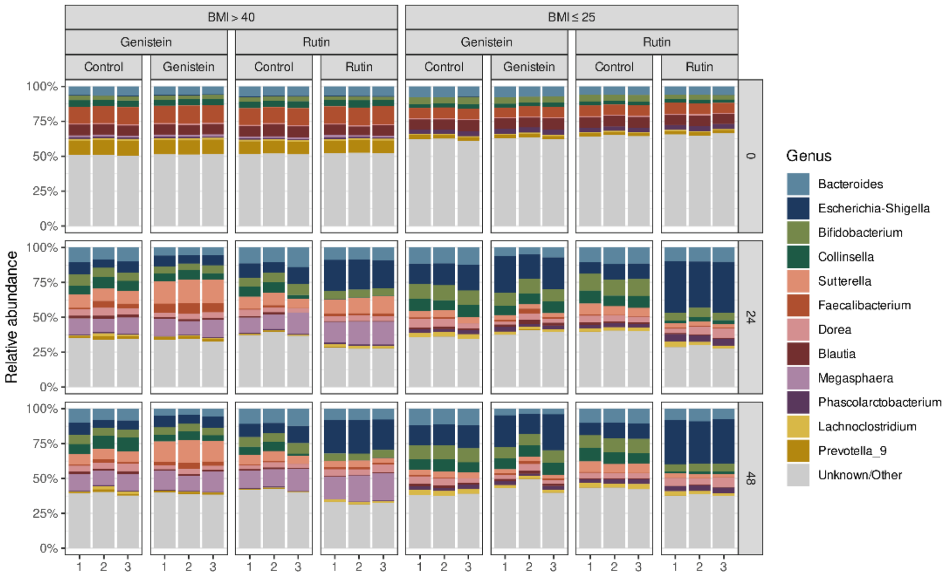


**Fig. S1 Time and treatment dependent changes in the composition of the bacterial communities** Changes in the compositions are shown as relative abundances for the most abundant representatives. On the horizontal axis separated in the two different pools BMI >40 and BMI <25 and the two different treatments rutin and genistein with the corresponding controls without the plant compounds for each of the three experimental replicates (1-3). On the vertical axis the separation for the three sampling times 0/ 24/ 48h. **A** composition plot at the level of the class, **B** at the level of the order, **C** at the level of the family, and **D** at the level of the genera.
